## Supplementary material for "Induced responses contribute to rapid plant adaptation to herbivory"

#### Supplementary figures

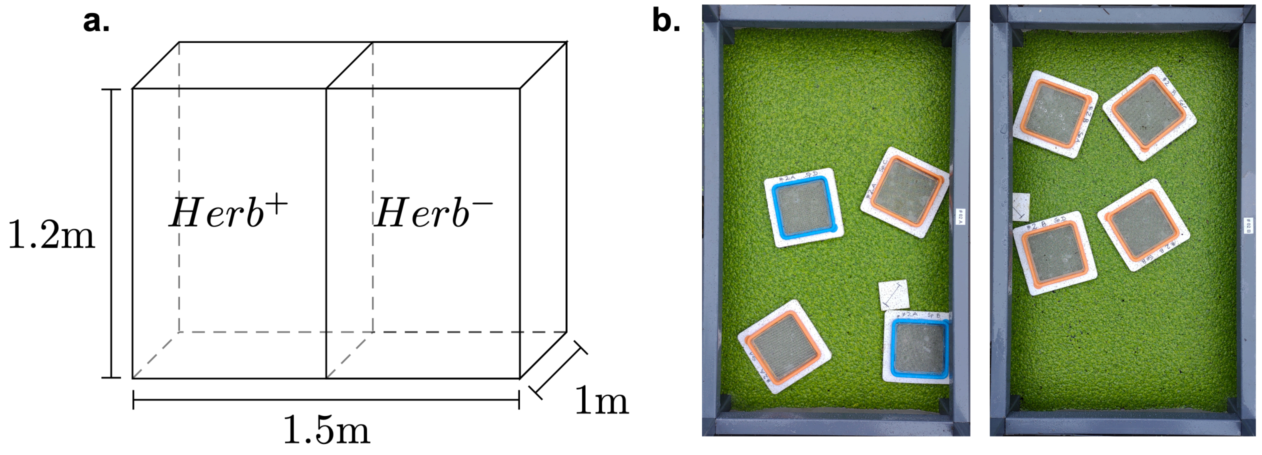

**Figure S1**. (a) Schematic representation of each pond. (b) Example of control (left) and herbivory (right) cages showing the floating boxes.

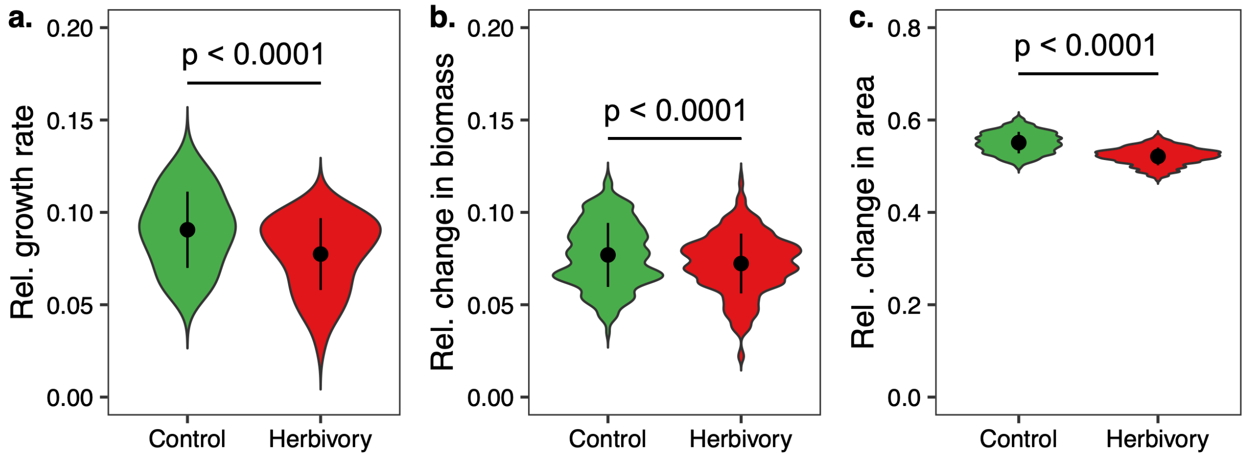

**Figure S2**. Relative growth rate (**a**; F = 36.72; df = 1; $p < 0.0001$), and change in biomass (**b**; F = 972.74; df = 1; $p < 0.0001$) and surface area (**c**; F = 28132; df = 1; $p < 0.0001$) within 2 weeks for plants exposed to 8 weeks of multigenerational herbivory or control conditions. Values for plots (**b**) and (**c**) are calculated by permutation ($n = 999$) with a set of reference fronds collected before the bioassay. For each group, dots represent mean values and bars show the standard deviation.

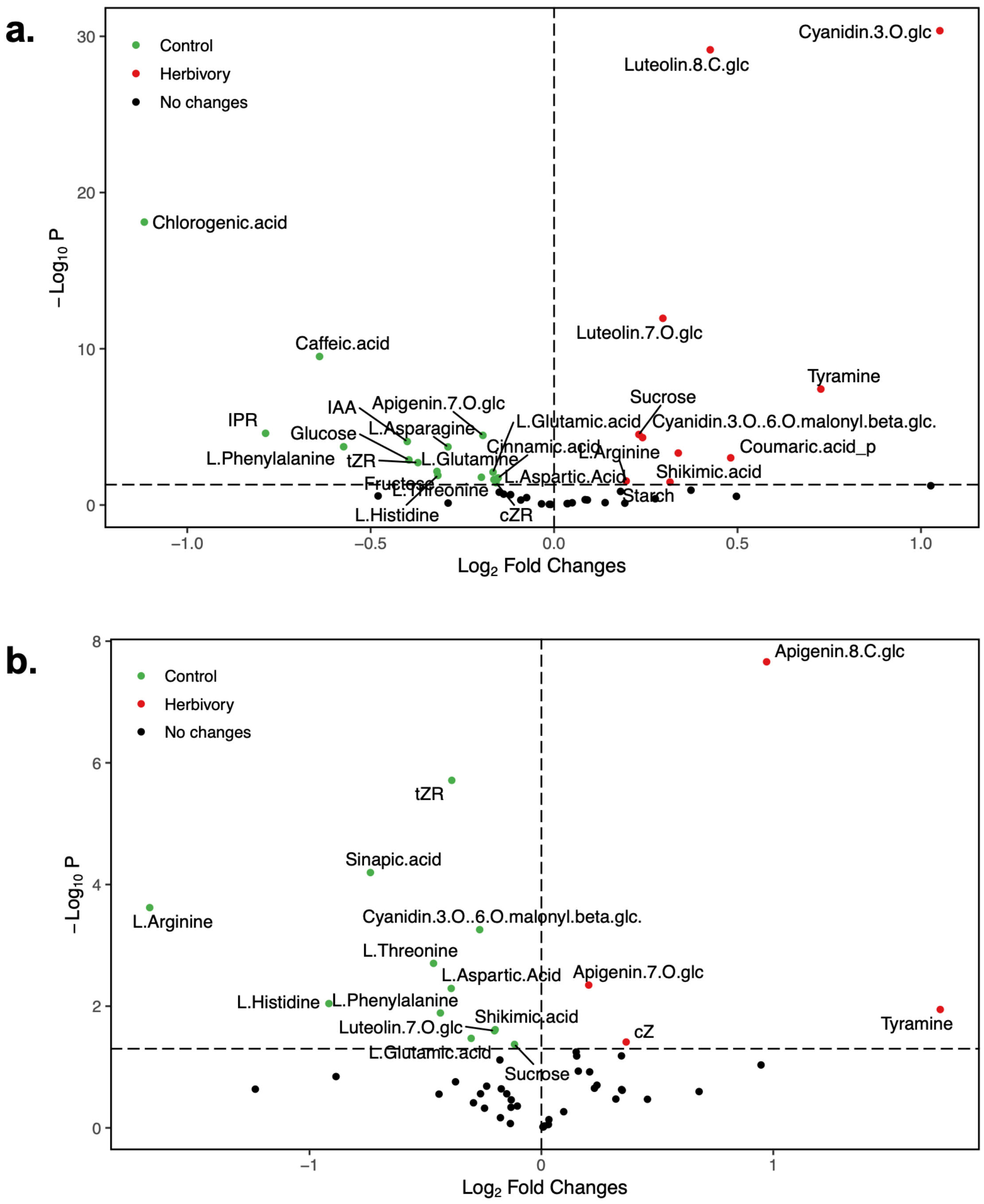

**Figure S3**. Volcano plot showing the metabolites of mixed popualtions with higher abundance in either control (left) or herbivory (right) treatments for first (**a**; week 12) and second (**b**; week 58) growing season.

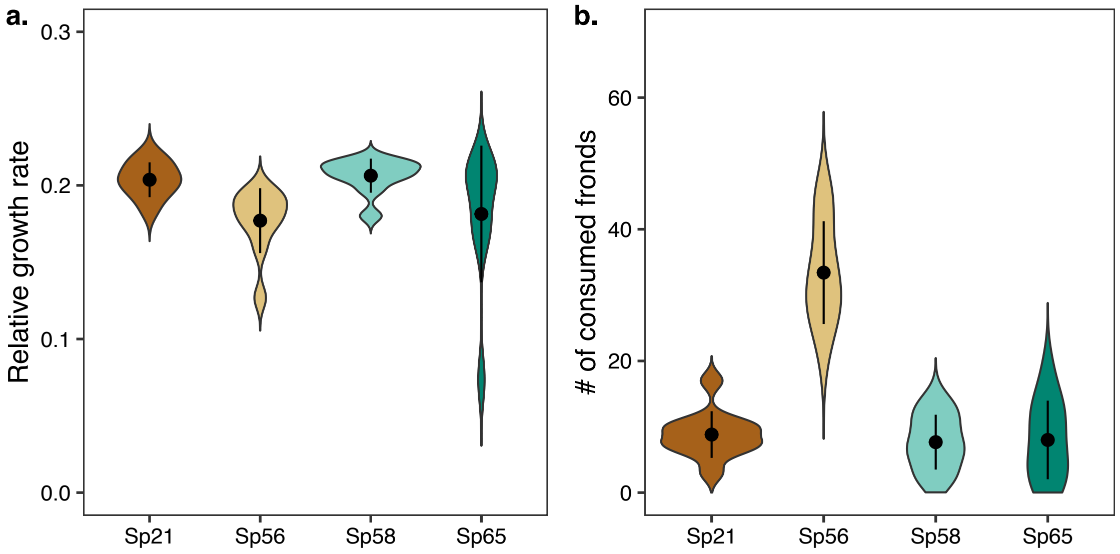

**Figure S4**. Relative intrinsic growth rate of each Spirodela polyrhiza genotype grown separatly under sterile and controlled conditions indoors (**a**; $F_{3,\,35} = 3.51$, $p = 0.02$), and number of consumed fronds for each of them after being exposed to 6 h of herbivory by a single snail (**b**; $F_{3,\,44} = 58.62$, $p < 0.0001$). For each group, dots represent mean values and bars show the standard deviation. **Methods**: To compare intrinsic growth rates and resistance of our four *Spirodela polyrhiza* genotypes, we grew gnotobiotic fronds of all genotypes separately in controlled conditions at 23 °C. Gnotobiotic plants were pre-cultivated under sterile conditions in full N-Medium for four weeks, before being transferred for pre-adaptation to 1:10 N-Medium under sterile conditions for another week. We then started the bioassays adding 30 gnotobiotic plants, separately for each genotype ($n$=10 boxes for Sp21, Sp56, Sp58, $n$=8 boxes of Sp65), to an autoclaved plastic box with a fine metal net on top of the lid to allow gas exchange and filled with 800 mL of fresh sterile 1:10 N-Medium. First, the 30 starting fronds were allowed to propagate for 13 days before harvested, and resulting fronds were counted. Second, we transferred 50 fronds from each box to a plastic filled with 200 mL of 1:10 N-Medium, we added one snail (from our laboratory culture, starved in 1:10 N-Medium for 24 prior) to each cup. Snails were removed after 6h, and we counted the number of consumed fronds.

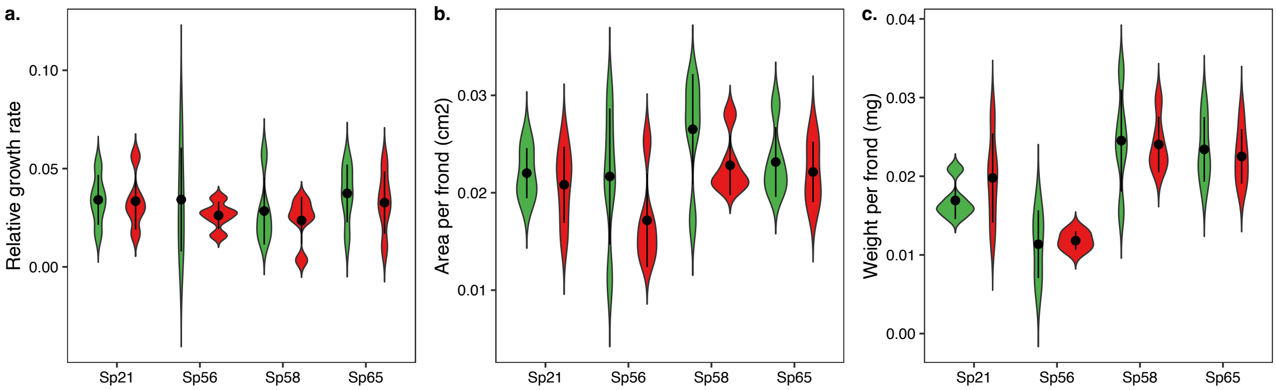

**Figure S5**. Relative growth rate (**a**), area per frond (**b**), and dry weight per frond (**c**) after a 13-days growth assay of individual *Spirodela polyrhiza* genotypes grown for 8 weeks in presence (red) or absence (green) of herbivores in outdoor ponds. Results from data analysis are reported in Tab. S5 below.

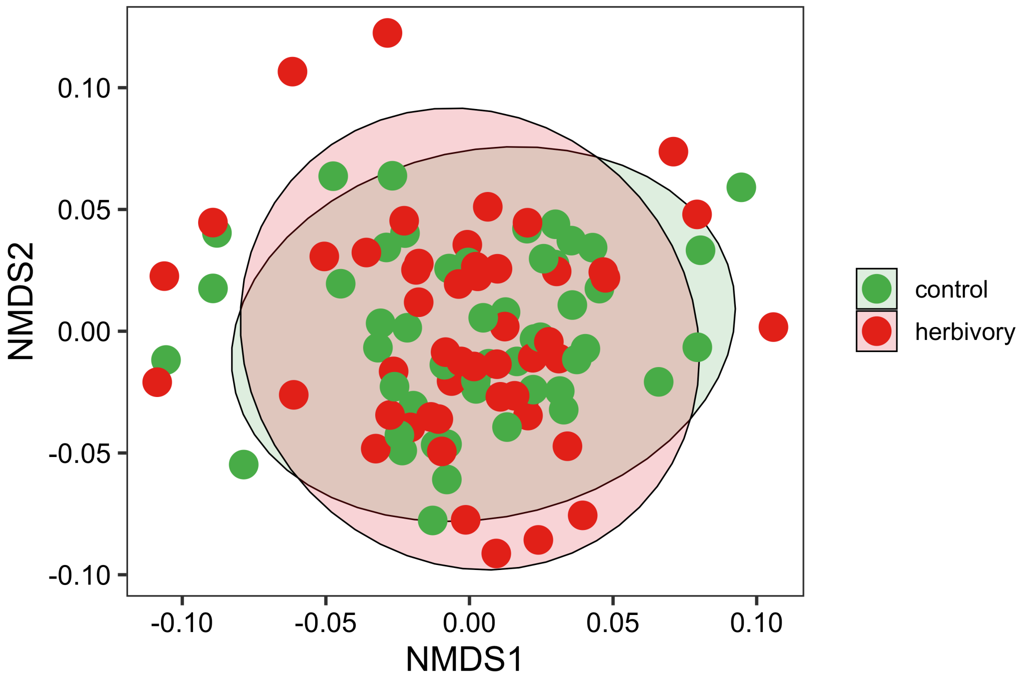

**Figure S6**. NMDS showing the overlap between water nutrient content in control and herbivory cages in outdoor ponds (data collected at weeks 4, 6, 8, 10, and 12). PERMANOVA analysis confirmed no differences between the two treatments ($F_{1,\,98} = 0.38$, $p = 0.41$; 999 permutations stratified over "pond" and "sampling timepoint").

#### Supplementary tables

**Table S1**. Geographical origins and Unified ID for each duckweed genotype used in our study.

| Genotype ID | Geographical origin | Unified ID |
|---|---|---|
| *Sp21* | Porstendorf, Germany | 9500 |
| *Sp56* | Pukkila, Finland | 9256 |
| *Sp58* | Szarvas, Hungary | 9560 |
| *Sp65* | Delta del Po, Veneto, Italy | 9413 |

**Table S2**. Linear mixed-effect models testing the influence of length of time of experimental evolution and treatment (herbivory/control) on *Spirodela polyrhiza* frond area and biomass.

| | | Area per frond | | Weight per frond | |
|---|---|---|---|---|---|
| **Factor** | **df** | **F** | **p** | **F** | **p** |
| *Timepoint* | 1 | 8.05 | 0.0052 | 2.51 | 0.1158 |
| *Treatment* | 1 | 103.19 | <0.0001 | 7.10 | 0.0086 |
| *Timepoint × Treatment* | 1 | 0.39 | 0.5324 | 0.61 | 0.4338 |

**Table S3**. Linear mixed-effect model testing the effect of treatment (herbivory/control) and genotype (Sp21, Sp56, Sp58, and Sp65) on the relative frequency of each *Spirodela polyrhiza* genotype within the experimental ponds after 12 weeks of experimental evolution approach in outdoor ponds. Both the pond ID and the sampling week were used as random factors.

| Factor | df | F | p |
|---|---|---|---|
| *Treatment* | 1 | 2.08 | 0.15 |
| *Genotype* | 3 | 52.25 | <0.0001 |
| *Treatment × Genotype* | 3 | 9.23 | <0.0001 |

**Table S4**. Linear mixed-effect models testing the influence of genotype and treatment (herbivory/control) on *Spirodela polyrhiza* frond area and biomass on individual genotypes grown in greenhouse for 8 weeks under herbivory or control conditions.

| | | Relative growth rate | | Area per frond | | Weight per frond | |
|---|---|---|---|---|---|---|---|
| **Factor** | **df** | **F** | **p** | **F** | **p** | **F** | **p** |
| *Genotype* | 3 | 0.87 | 0.46 | 2.63 | 0.07 | 22.81 | <0.0001 |
| *Treatment* | 1 | 1.11 | 0.30 | 3.68 | 0.06 | 0.16 | 0.69 |
| *Timepoint × Treatment* | 3 | 0.12 | 0.94 | 1.42 | 0.73 | 0.49 | 0.68 |

**Table S5**. PERMANOVA testing the effect of treatment (herbivory/control) and length of experimental evolution (week) on the structure of bacterial, fungal, and algal communities associated with *Spirodela polyrhiza*.

| | | Bacteria and fungi | | | Algae | | |
|---|---|---|---|---|---|---|---|
| **Factor** | **df** | $R^2$ | **F** | **p** | $R^2$ | **F** | **p** |
| *Treatment* | 1 | 0.05 | 2.54 | 0.007 | 0.06 | 4.41 | 0.036 |
| *Week* | 1 | 0.15 | 7.16 | 0.001 | 0.34 | 22.54 | 0.001 |
| *Treatment × Week* | 1 | 0.02 | 1.04 | 0.186 | 0.04 | 3.13 | 0.070 |

**Table S6**. Linear mixed-effect models testing the effect of genotype (Sp21, Sp56, Sp58, and Sp65) and treatment (herbivory/control) on the growth (changes in biomass and area) and resistance (consumed biomass and area) of single genotype of *Spirodela polyrhiza* grown in the experimental ponds.

| | | Change in biomass | | Change in area | | Consumed biomass | | Consumed area | |
|---|---|---|---|---|---|---|---|---|---|
| **Factor** | **df** | **F** | **p** | **F** | **p** | **F** | **p** | **F** | **p** |
| *Genotype* | 3 | 2.83 | 0.04 | 0.11 | 0.95 | 137.94 | <0.001 | 24.25 | <0.001 |
| *Treatment* | 1 | 0.26 | 0.61 | 0.37 | 0.54 | 41.95 | <0.001 | 4.20 | 0.04 |
| *Genotype × Treatment* | 3 | 0.23 | 0.87 | 0.34 | 0.79 | 78.71 | <0.001 | 32.57 | <0.001 |

**Table S7**. Linear mixed-effect model testing the effect of treatment (herbivory/control) on the number of consumed fronds of single genotypes after 24h of herbivory for each *Spirodela polyrhiza* genotype, grown for 8 weeks outdoors in cages with or without herbivores, but protected from snails using fine nets.

| **Genotype** | **F** | **df** | **p** |
|---|---|---|---|
| *Sp21* | 0.001 | 1 | 0.96 |
| *Sp56* | 0.037 | 1 | 0.84 |
| *Sp58* | 2.458 | 1 | 0.11 |
| *Sp65* | 3.669 | 1 | 0.05 |

**Table S8**. Taxa differentially abundant between plants (genotypes Sp21 and Sp65) grown under control or herbivory-induced microbiota environment.

| Genotype | Marker | Treatment with higher abundance | Genus | Number of ASVs |
|---|---|---|---|---|
| Sp21 | 16S | Herbivory | *Cetobacterium* | 3 |
| | | Control | - | - |
| Sp65 | 16S | Herbivory | *Bacteroides* | 1 |
| | | Control | *Ralstonia* | 1 |
| | | | Unid. Muribaculaceae | 3 |
| | | | *Clostridium* | 1 |
| | | | Unid. Lachnospiraceae | 1 |
| | | | *Cetobacterium* | 2 |

**Table S9**. Linear-mixed effects models testing the influence of herbivory on water nutrient levels (and pH) during the first growing season. Both the pond ID and the sampling timepoints were used as random factors.

| Nutrient | Control (mean ± SD) | Herbivore (mean ± SD) | df | F | p |
|---|---|---|---|---|---|
| *Cl* | 25.90 ± 4.44 | 25.27 ± 4.96 | 1 | 4.77 | 0.03 |
| $NO_2$ | 4.68 ± 1.24 | 4.63 ± 1.12 | 1 | 0.19 | 0.66 |
| $NO_3$ | 8.69 ± 5.50 | 8.28 ± 5.21 | 1 | 2.66 | 0.10 |
| $PO_4$ | 1.67 ± 1.16 | 1.73 ± 1.06 | 1 | 0.25 | 0.61 |
| $SO_4$ | 11.98 ± 4.73 | 11.58 ± 4.66 | 1 | 4.42 | 0.03 |
| *Na* | 25.48 ± 3.21 | 24.98 ± 3.73 | 1 | 2.22 | 0.14 |
| $NH_4$ | 1.95 ± 2.84 | 1.91 ± 2.76 | 1 | 0.35 | 0.55 |
| *K* | 35.37 ± 7.86 | 34.72 ± 8.63 | 1 | 0.67 | 0.41 |
| *Ca* | 33.44 ± 6.30 | 32.87 ± 6.35 | 1 | 0.81 | 0.37 |
| *Mg* | 4.44 ± 0.61 | 4.37 ± 0.68 | 1 | 1.37 | 0.24 |
| *pH* | 7.49 ± 0.48 | 7.50 ± 0.48 | 1 | 0.36 | 0.54 |

#### Supplementary methods - water nutrients analysis

While our treatment cages shared the same water body of a pond, thus per se allowing exchange of nutrients, we tested individual nutrient levels across treatments and ponds using ion chromatography (792 basic IC, 732 detector, Metrohm, Herisau, Switzerland). Samples were taken every two weeks (from experimental setup to week 12) during the first season (n=10 per treatment cage) and once during the second growth season at week 58 (n=7 per treatment cage). We sampled 10 mL per cage for anion and cation analysis, respectively. When collecting samples from each cage, we pooled water from five different locations at a depth of 10 cm below the water surface. Anion samples were measured right away, while pH of cation samples was adjusted to pH 3 using 25 % $HNO_3$ before storage at -20°C until analysis. Anions (Cl, $NO_2$, $NO_3$, $PO_4$, $SO_4$) were measured using: column Metrosep A Supp 5 150, 4.0 x 150 mm, eluent 3.2 mM $Na_2CO_3$ and 1 mM $NaHCO_3$, flow rate 0.7 ml/min, injection volume 20 µl. Cations (Na, NH4, K, Ca, Na) were measured using: column Metrosep C2 250, 4.0 x 250 mm, eluent 1.7 mmol/L nitric acid, 0.7 mmol/L dipicolinic acid, flow rate 0.9 ml/min, injection volume 20 µl. Quantification was based on external standard ion mixture using a 6-point-reference curve (0.5, 5, 10, 25, 50, 100 mg/L for anions and 0.5, 5, 10, 15, 25, 50 m/L for cations). Additionally, throughout the growth seasons we measured the pH of each pond using a Cond3310, (WTW, Xylem Analytics) instrument.

The effect of herbivory on water nutrients and pH was tested by fitting a linear mixed effects model specifying treatment (herbivory/control) as fixed factor, and pond and timepoint as random effects, using the formula: $\sim$ treatment * (1|timepoint) * (1|pond). In addition, we tested the effects of overall nutrient content through PERMANOVA on data collected at week 4, 6, 8, 10, and 12 (999 permutations, stratified over “pond” and “sampling timepoint”).

### Supplementary methods - metabolomics

Extraction and analysis of diverse primary and secondary metabolites was done similar as described by Schäfer et al. [1]. A detailed list of the analyzed compound and the respective analytical methods can be found in Table S10-16. In brief, approximately 10 mg fine ground, freeze dried plant tissue was extracted with acidified methanol containing the following internal standards: 10 ng $D_6$-ABA (OlChemIm); 10 ng $D_5$-JA (Sigma-Aldrich); 10 ng $D_4$-SA (OlChemIm); 0.5 ng $D_5$-tZ (OlChemIm); 0.1 ng $D_6$-IP (OlChemIm); 0.1 ng $D_6$-IPR (OlChemIm); 1 ng $^{13}C_6$-IAA (OlChemIm); 3 ng 4-methylumbelliferone (4-MU) (Sigma-Aldrich). For the analysis of different high abundant compounds, a small aliquot (A1) of the extract was taken aside, while the rest of the extract (E1) was used for further processing. High abundant compounds, such as amino acids and related compounds were analyzed in a 1:100 dilution of A1 in an aqueous mix of isotope-labeled amino acids (algal amino acid mixture-$^{13}$C-$^{15}$N; Sigma-Aldrich; Method 1A and 1B). Soluble sugars were measured in a 1:125 dilution of A1 in 70% methanol containing sorbitol as internal standard (Fluka Biochemika; Method 1C). The major secondary metabolites were analyzed in the rest of the aliquot (A1) without further dilution (Method 1D). After a second extraction of the plant tissue with acidified methanol this supernatant was combined with the first extract (E1) and subsequently purified on a reversed phase (RP) solid phase extraction (SPE) column (HR-X, Macherey-Nagel) and a mixed-mode RP-cation exchange SPE column (HR-XC, Macherey-Nagel). The compounds were bound to the HR-XC column, different washing steps were applied (e.g. to remove CK-phosphates) and the analytes of interest were stepwise eluted from the column to generate a fraction containing the acidic phytohormones, such as JA, ABA and IAA, as well as a fraction containing the CKs. The eluted fractions were analyzed directly (Method 2A) or after an additional concentration step (Method 2B) and (Method 3) dependent on the abundance and instrument sensitivity for the respective compounds. Starch content was determined from the remaining material (pellet) after the two rounds of extraction with acidified methanol. For this, the material was subjected to two additional washing steps with 50% ethanol at 80 °C to remove soluble sugars. Subsequently, a sub-fraction of the material was resuspended in water and incubated at 98 °C to gelatinize the starch, which was then digested to glucose by amyloglucosidase (from Aspergillus niger, Roche) and ¸ -amylase (from porcine pancreas, Sigma) at 37 °C. The glucose content was then measured in a 1:250 dilution of this solution in 70% methanol containing sorbitol as internal standard the same way as for the soluble sugars before (Method 1C).

The major secondary metabolites (Method 1D) were analyzed by HPLC-PDA. Chromatographic separation was done on a Shimadzu Nexera XR LC-System equipped with an EC 4/3 Nucleodur Sphinx RP pre-column (5 µm, Macherey-Nagel) and a Nucleodur Sphinx RP column (250x4.6 mm, 5µm, Macherey-Nagel). The mobile phase comprised 0.2% formic acid (Fisher Chemical), 0.1% acetonitrile (Fisher Chemical) in water as Solvent A and acetonitrile (Fisher Chemical) as Solvent B. The column oven was set to 20 °C and the flow rate to 1300 µL/min. Measurement was performed with a PDA detector.

Other compounds (Method 1A, 1B, 1C, 2A, 2B and 3) were analyzed by LC-MS/MS. Chromatographic separation for was done on a Shimadzu Nexera X3 LC-System. For method 1A, 1B, 2A, 2B and 3 the system was equipped with an Agilent 1290 infinity II inline filter (0,3 µm) and a ZORBAX RRHD Eclipse XDB-C18 column (3x50 mm, 1.8 µm; Agilent Technologies). The mobile phase comprised 0.05% formic acid (Fisher Chemical), 0.1% acetonitrile (Fisher Chemical) in water as Solvent A and methanol (Fisher Chemical) as Solvent B. The column oven was set to 42 °C and the flow rate to 500 µL/min. For method 1C the system was equipped with an Agilent 1290 infinity II inline filter (0,3 µm) and an apHera NH2 column (150x4.6 mm, 5 µm; Supelco). The mobile phase comprised 0.1% acetonitrile (Fisher Chemical) in water as Solvent A and acetonitrile (Fisher Chemical) as Solvent B. The column oven was set to 25 °C and the flow rate to 700 µL/min. The measurements were performed on a Shimadzu LCMS-8060 equipped with an ESI source, which was operated in multi-reaction-monitoring (MRM) modus. Settings were as follows: Nebulizing Gas Flow: 3 L/min; Heating Gas Flow: 10 L/min; Drying Gas Flow: 10 L/min; Interface Temperature: 300 °C; DL Temperature

250 °C; Heat Block Temperature: 400 °C; CID Gas: 270 kPa; Q1 Resolution: Unit; Q3 Resolution: Unit.

The gradient programs for all methods are given in Table S17-23 and the detector settings (MRM-settings or absorption wavelength, respectively) in Table S10-16.

**Table S10**. MRM-settings and retention times of analytes of Method 1A.

| **Analyte** | **RT [min]** | **Q1 [m/z]** | **Q3 [m/z]** | **Dwell time [ms]** | **CE [V]** | **Q1/Q3 Pre Bias [V]** | **ISTD** [a] |
|---|---|---|---|---|---|---|---|
| Ala | 0.450 | (+)90.05 | 44.20 | 20 | -13 | -10/-18 | $^{13}C_3,^{15}N_1$-Ala |
| Arg | 0.420 | (+)175.12 | 60.20 | 20 | -14 | -20/-24 | $^{13}C_6,^{15}N_4$-Arg |
| Asp | 0.450 | (+)134.04 | 88.20 | 20 | -12 | -13/-16 | $^{13}C_4,^{15}N_n$-$Asx_{Asp}$ [b] |
| Glu | 0.450 | (+)148.06 | 102.15 | 20 | -13 | -10/-19 | $^{13}C_5,^{15}N_n$-$Glx_{Glu}$ [b] |
| Pro | 0.485 | (+)116.07 | 70.20 | 20 | -17 | -11/-20 | $^{13}C_5,^{15}N_1$-Pro |
| Ser | 0.435 | (+)106.05 | 60.20 | 20 | -14 | -10/-23 | $^{13}C_3,^{15}N_1$-Ser |
| Thr | 0.450 | (+)120.07 | 74.20 | 20 | -12 | -12/-30 | $^{13}C_4,^{15}N_1$-Thr |
| Asn | 0.435 | (+)133.06 | 87.20 | 20 | -11 | -13/-16 | $^{13}C_4,^{15}N_n$-$Asx_{Asn}$ [b] |
| Gln | 0.450 | (+)147.08 | 130.15 | 20 | -15 | -10/-13 | $^{13}C_5,^{15}N_n$-$Glx_{Gln}$ [b] |
| Val | 0.627 | (+)118.09 | 72.20 | 20 | -12 | -22/-28 | $^{13}C_5,^{15}N_1$-Val |
| Ile | 1.165 | (+)132.10 | 86.15 | 122 | -11 | -20/-20 | $^{13}C_6,^{15}N_1$-Ile |
| Leu | 1.255 | (+)132.10 | 86.15 | 122 | -11 | -20/-20 | $^{13}C_6,^{15}N_1$-Leu |
| Phe | 2.520 | (+)166.09 | 120.20 | 247 | -15 | -20/-20 | $^{13}C_9,^{15}N_1$-Phe |
| Trp | 3.260 | (+)205.10 | 188.10 | 32 | -10 | -20/-20 | $^{13}C_9,^{15}N_1$-Phe (1.76) |
| | | (+)205.10 | 118.20 | 32 | -26 | -20/-20 | |
| $^{13}C_3$, $^{15}N_1$-Ala | 0.450 | (+)94.06 | 47.20 | 20 | -13 | -10/-18 | |
| $^{13}C_6$, $^{15}N_4$-Arg | 0.438 | (+)185.13 | 64.15 | 20 | -14 | -20/-24 | |
| $^{13}C_4$, $^{15}N_n$-Asx | 0.450 | (+)139.06 | 92.25 | 20 | -12 | -13/-16 | |
| $^{13}C_5$, $^{15}N_n$-Glx | 0.450 | (+)154.07 | 107.20 | 20 | -13 | -10/-19 | |
| $^{13}C_5$, $^{15}N_1$-Pro | 0.485 | (+)122.08 | 75.20 | 20 | -17 | -11/-20 | |
| $^{13}C_3$, $^{15}N_1$-Ser | 0.435 | (+)110.06 | 63.20 | 20 | -14 | -10/-23 | |
| $^{13}C_4$, $^{15}N_1$-Thr | 0.450 | (+)125.08 | 78.20 | 20 | -12 | -12/-30 | |
| $^{13}C_5$, $^{15}N_1$-Val | 0.627 | (+)124.10 | 77.20 | 20 | -12 | -22/-28 | |
| $^{13}C_6$, $^{15}N_1$-Ile | 1.165 | (+)139.12 | 92.25 | 122 | -11 | -20/-20 | |
| $^{13}C_6$, $^{15}N_1$-Leu | 1.255 | (+)139.12 | 92.25 | 122 | -11 | -20/-20 | |
| $^{13}C_9$, $^{15}N_1$-Phe | 2.520 | (+)176.11 | 129.25 | 247 | -15 | -20/-20 | |

RT: retention time
CE: collision energy
ISTD: internal standard
Qualifiers are highlighted in grey
[a] Incl. matrix corrected response factor ($n_{Analyte} = x * n_{ISTD}$) in brackets
[b] Asx and Glx handled as Asn, Asp, Glu or Gln equivalents, respectively

**Table S11**. MRM-settings and retention times of analytes of Method 1B.

| Analyte | RT [min] | Q1 [m/z] | Q3 [m/z] | Dwell time [ms] | CE [V] | Q1/Q3 Pre Bias [V] | ISTD[a] |
|---|---|---|---|---|---|---|---|
| Gly | 0.46 | (+)76.04 | 30.20 | 50 | -12 | -14/-11 | $^{13}C_2$,$^{15}N_1$-Gly |
| His | 0.44 | (+)156.08 | 110.20 | 50 | -15 | -30/-21 | $^{13}C_6$,$^{15}N_3$-His |
| Met | 0.75 | (+)150.06 | 61.15 | 50 | -22 | -10/-24 | $^{13}C_5$,$^{15}N_1$-Met |
| Shikimic acid | 0.63 | (-)173.05 | 93.10 | 23 | 15 | 12/21 | $^{13}C_9$,$^{15}N_1$-Tyr (37.77) |
| | | (-)173.05 | 111.20 | 23 | 13 | 12/10 | |
| Tyr | 1.13 | (+)182.08 | 136.20 | 50 | -15 | -18/-14 | $^{13}C_9$,$^{15}N_1$-Tyr |
| Tyramine | 1.16 | (+)138.09 | 121.15 | 23 | -15 | -20/-20 | $^{13}C_9$,$^{15}N_1$-Tyr (0.51) |
| | | (+)138.09 | 77.15 | 23 | -30 | -20/-20 | |
| Tryptamine | 3.28 | (+)161.15 | 144.15 | 297 | -15 | -20/-20 | $^{13}C_9$,$^{15}N_1$-Tyr (0.36) |
| | | (+)161.15 | 117.15 | 297 | -25 | -20/-22 | |
| Apigenin | 4.12 | (-)269.05 | 151.20 | 147 | 25 | 30/14 | $^{13}C_9$,$^{15}N_1$-Tyr (0.96) |
| | | (-)269.05 | 149.15 | 147 | 24 | 29/28 | |
| Luteolin | 4.025 | (-)285.04 | 132.20 | 147 | 48 | 30/26 | $^{13}C_9$,$^{15}N_1$-Tyr (3.44) |
| | | (-)285.04 | 175.20 | 147 | 26 | 19/11 | |
| $^{13}C_2$, $^{15}N_1$-Gly | 0.46 | (+)79.04 | 32.10 | 50 | -12 | -14/-11 | |
| $^{13}C_6$, $^{15}N_3$-His | 0.44 | (+)165.09 | 118.20 | 50 | -15 | -30/-21 | |
| $^{13}C_5$, $^{15}N_1$-Met | 0.75 | (+)156.07 | 63.15 | 50 | -22 | -10/-24 | |
| $^{13}C_9$, $^{15}N_1$-Tyr | 1.13 | (+)192.11 | 145.20 | 50 | -15 | -18/-14 | |

RT: retention time
CE: collision energy
ISTD: internal standard
Qualifiers are highlighted in grey
[a] Incl. matrix corrected response factor ($n_{Analyte} = x * n_{ISTD}$) in brackets

**Table S12**. MRM-settings and retention times of analytes of Method 1C.

| Analyte | RT [min] | Q1 [m/z] | Q3 [m/z] | Dwell time [ms] | CE [V] | Q1/Q3 Pre Bias [V] | ISTD[a] |
|---|---|---|---|---|---|---|---|
| Fructose | 7.8 | (-)179.06 | 88.90 | 19 | 9 | 12/13 | Sorbitol[b] |
| | | (-)179.06 | 59.00 | 19 | 17 | 12/11 | |
| | | (-)179.06 | 71.00 | 19 | 16 | 11/11 | |
| Glucose | 9.2 | (-)179.06 | 88.90 | 19 | 9 | 12/13 | Sorbitol[b] |
| | | (-)179.06 | 59.00 | 19 | 17 | 12/11 | |
| | | (-)179.06 | 71.00 | 19 | 16 | 11/11 | |
| Sucrose | 8.4 | (-)341.11 | 89.10 | 19 | 23 | 12/10 | Sorbitol[b] |
| | | (-)341.11 | 179.15 | 19 | 15 | 30/30 | |
| | | (-)341.11 | 59.05 | 19 | 36 | 29/23 | |
| Sorbitol | 10.8 | (-)181.07 | 88.85 | 19 | 16 | 11/17 | |
| | | (-)181.07 | 59.00 | 19 | 22 | 11/10 | |
| | | (-)181.07 | 71.00 | 19 | 22 | 30/29 | |

RT: retention time
CE: collision energy
ISTD: internal standard
Qualifiers are highlighted in grey
[a] Incl. matrix corrected response factor ($n_{Analyte} = x * n_{ISTD}$) in brackets
[b] Response factor calculated based on an external dilution curve run with each batch of samples

**Table S13**. Absorption and retention times of analytes of Method 1D. Quantification based on an external dilution curve with identical standards, except for the putative chlorogenic acid isomere that was based on the chlorogenic acid standard, and cyanidin-3-O-(6-O-malonyl-beta-glucoside) that was quantified based on the molar quantity of the cyanidin-3-O-glucoside standard.

| Analyte | RT [min] | Absortion wave lenght [nm] |
|---|---|---|
| Cyanidin-3-O-glucoside | 7.055 | 517 |
| Chlorogenic acid | 8.055 | 328 |
| Putative Chlorogenic acid isomere | 8.210 | 328 |
| Cyanidin-3-O-(6-O-malonyl-beta-glucoside) | 9.410 | 517 |
| Luteolin-8-C glucoside | 11.295 | 348 |
| Apigenin-8-C-glucoside | 12.495 | 337 |
| Luteolin-7-O-glucoside | 13.085 | 348 |
| Apigenin-7-O-glucoside | 14.325 | 337 |

RT: retention time

**Table S14**. MRM-settings and retention times of analytes of Method 2A

| Analyte | RT [min] | Q1 [m/z] | Q3 [m/z] | Dwell time [ms] | CE [V] | Q1/Q3 Pre Bias [V] | ISTD |
|---|---|---|---|---|---|---|---|
| SA | 2.57 | (-)137.02 | 92.95 | 20 | 19 | 27 / 19 | $D_4$-SA |
| | | (-)137.02 | 65.00 | 20 | 32 | 27 / 10 | |
| ABA | 2.71 | (-)263.13 | 153.10 | 20 | 13 | 18 / 29 | $D_6$-ABA |
| | | (-)263.13 | 204.15 | 20 | 20 | 18 / 12 | |
| JA | 3.20 | (-)209.12 | 59.00 | 20 | 14 | 14 / 11 | $D_5$-JA |
| | | (-)209.12 | 40.95 | 20 | 44 | 14 / 14 | |
| JA-Ile | 4.57 | (-)322.20 | 130.10 | 135 | 22 | 24 / 20 | $D_5$-JA (0.084) |
| | | (-)322.20 | 128.15 | 135 | 22 | 24 / 26 | |
| OPDA | 6.21 | (-)291.20 | 165.25 | 135 | 22 | 20 / 10 | $D_5$-JA[b] |
| | | (-)291.20 | 247.15 | 135 | 20 | 20 / 23 | |
| $D_4$-SA | 2.56 | (-)141.05 | 97.00 | 20 | 19 | 27 / 15 | |
| | | (-)141.05 | 69.05 | 20 | 32 | 27 / 17 | |
| $D_6$-ABA | 2.70 | (-)269.17 | 159.20 | 20 | 13 | 18 / 30 | |
| | | (-)269.17 | 207.20 | 20 | 20 | 18 / 20 | |
| $D_5$-JA | 3.19 | (-)214.15 | 62.00 | 20 | 14 | 14/ 10 | |
| | | (-)214.15 | 42.00 | 20 | 44 | 14 / 15 | |

RT: retention time
CE: collision energy
ISTD: internal standard
Qualifiers are highlighted in grey
[a] Incl. matrix corrected response factor ($n_{Analyte} = x * n_{ISTD}$) in brackets
[b] Relative quantification

**Table S15**.MRM-settings and retention times of analytes of Method 2B.

| Analyte | RT [min] | Q1 [m/z] | Q3 [m/z] | Dwell time [ms] | CE [V] | Q1/Q3 Pre Bias [V] | ISTD[a] |
|---|---|---|---|---|---|---|---|
| Caffeic acid | 2.275 | (-) 179.03 | 135.05 | 20 | 18 | 12 / 26 | 4-MU[b] |
| | | (-) 179.03 | 134.15 | 20 | 25 | 12 / 12 | |
| | | (-) 179.03 | 107.10 | 20 | 24 | 19 / 21 | |
| *p*-Coumaric acid | 2.624 | (-) 163.04 | 119.10 | 20 | 17 | 11 / 11 | 4-MU[b] |
| | | (-) 163.04 | 93.05 | 20 | 32 | 11 / 19 | |
| Sinapic acid | 2.76 | (+)225.08 | 207.15 | 20 | -9 | -15 / -23 | 4-MUb |
| | | (+)225.08 | 119.20 | 20 | -19 | -15 / -12 | |
| IAA | 3.237 | (+)176.07 | 130.10 | 20 | -16 | -12 / -23 | $^{13}C_6$-IAA (0.42) |
| | | (+)176.07 | 130.20 | 20 | -31 | -12 / -10 | |
| Cinamic acid | 5.002 | (-) 147.05 | 103.10 | 100 | 15 | 17 / 20 | 4-MU[b] |
| | | (-) 147.05 | 77.10 | 100 | 23 | 10 / 18 | |
| $^{13}C_6$-IAA | 3.237 | (+)182.09 | 109.20 | 20 | -31 | -12 / -19 | |
| | | (+)182.09 | 136.25 | 20 | -16 | -12 / -13 | |
| 4-MU | 3.344 | (+)177.05 | 77.20 | 20 | -35 | -12 / -30 | |
| | | (+)177.05 | 105.20 | 20 | -21 | -12 / -20 | |

RT: retention time
CE: collision energy
ISTD: internal standard
Qualifiers are highlighted in grey
[a] Incl. matrix corrected response factor ($n_{Analyte} = x * n_{ISTD}$) in brackets
[b] Relative quantification

**Table S16**. MRM-settings and retention times of analytes of Method 3.

| Analyte | RT [min] | Q1 [m/z] | Q3 [m/z] | Dwell time [ms] | CE [V] | Q1/Q3 Pre Bias [V] | ISTD[a] |
|---|---|---|---|---|---|---|---|
| tZ | 2.527 | (+)220.12 | 136.10 | 47 | -20 | -20 / -20 | $D_5$-tZ |
| | | (+)220.12 | 119.15 | 47 | -33 | -22 / -12 | |
| cZ | 2.83 | (+)220.12 | 136.10 | 47 | -20 | -20 / -20 | $D_5$-tZ (0.94) |
| | | (+)220.12 | 119.15 | 47 | -33 | -22 / -12 | |
| tZR | 4.292 | (+)352.16 | 220.15 | 30 | -20 | -20 / -20 | $D_5$-tZ (0.17) |
| | | (+)352.16 | 136.15 | 30 | -30 | -20 / -20 | |
| cZR | 4.696 | (+)352.16 | 220.15 | 30 | -20 | -20 / -20 | $D_5$-tZ (0.17) |
| | | (+)352.16 | 136.15 | 30 | -33 | -20 / -20 | |
| IP | 5.648 | (+)204.12 | 136.10 | 30 | -15 | -20 / -20 | $D_6$-IP |
| | | (+)204.12 | 119.10 | 30 | -30 | -20 / -20 | |
| IPR | 6.39 | (+)336.17 | 204.20 | 47 | -20 | -20 / -20 | $D_6$-IPR |
| | | (+)336.17 | 136.10 | 47 | -30 | -20 / -20 | |
| $D_5$-tZ | 2.52 | (+)225.15 | 137.20 | 47 | -20 | -21 / -22 | |
| | | (+)225.15 | 119.20 | 47 | -36 | -15 / -12 | |
| $D_6$-IP | 5.511 | (+)210.16 | 137.20 | 30 | -15 | -20 / -20 | |
| | | (+)210.16 | 119.15 | 30 | -33 | -14 / -22 | |
| $D_6$-IPR | 6.363 | (+)342.20 | 210.25 | 47 | -20 | -20 / -20 | |
| | | (+)342.20 | 137.15 | 47 | -30 | -20 / -20 | |

RT: retention time
CE: collision energy
ISTD: internal standard
Qualifiers are highlighted in grey
[a] Incl. matrix corrected response factor ($n_{Analyte} = x * n_{ISTD}$) in brackets

**Table S17**. Solvent settings used for Method 1A.

| Time [min] | B [%] |
|---|---|
| 0.0 | 2 |
| 1.5 | 2 |
| 3.5 | 100 |
| 4.5 | 100 |
| 5.0 | 2 |
| 6.0 | 2 |

**Table S18**. Solvent settings used for Method 1B.

| Time [min] | B [%] |
|---|---|
| 0.0 | 2 |
| 1.5 | 2 |
| 3.5 | 100 |
| 4.5 | 100 |
| 5.0 | 2 |
| 6.0 | 2 |

**Table S19**. Solvent settings used for Method 1C.

| Time [min] | B [%] |
|---|---|
| 0 | 80 |
| 13 | 60 |
| 14 | 80 |
| 20 | 80 |

**Table S20**. Solvent settings used for Method 1D.

| Time [min] | B [%] |
|---|---|
| 0.0 | 10 |
| 8.0 | 21 |
| 18.0 | 49 |
| 18.1 | 100 |
| 19.0 | 100 |
| 19.1 | 10 |
| 24.0 | 10 |

**Table S21**. Solvent settings used for Method 2A.

| **Time [min]** | **B [%]** |
|---|---|
| 0.0 | 10 |
| 0.5 | 10 |
| 1.0 | 55 |
| 4.5 | 65 |
| 5.5 | 100 |
| 6.5 | 100 |
| 7.0 | 10 |
| 8.0 | 10 |

**Table S22**. Solvent settings used for Method 2B.

| **Time [min]** | **B [%]** |
|---|---|
| 0.0 | 10 |
| 0.5 | 10 |
| 1.0 | 39 |
| 3.5 | 41 |
| 3.6 | 50 |
| 8.0 | 60 |
| 8.5 | 100 |
| 9.5 | 100 |
| 10.0 | 10 |
| 11.0 | 10 |

**Table S23**. Solvent settings used for Method 3.

| **Time [min]** | **B [%]** |
|---|---|
| 0.0 | 5 |
| 0.5 | 5 |
| 0.7 | 15 |
| 3.5 | 25 |
| 6.5 | 70 |
| 6.7 | 100 |
| 7.7 | 100 |
| 8.0 | 5 |
| 9.0 | 5 |
